# Initial neural representation of individual rules after first-time instruction

**DOI:** 10.1101/598276

**Authors:** Hannes Ruge, Theo A. J. Schäfer, Katharina Zwosta, Holger Mohr, Uta Wolfensteller

**Affiliations:** Technische Universität Dresden, Germany; Max-Planck-Institute for Human Cognitive and Brain Sciences, Leipzig, Germany

## Abstract

By following explicit instructions humans can instantaneously get the hang of tasks they have never performed before. Here, we used a specially calibrated multivariate analysis technique to uncover the elusive representational states following newly instructed arbitrary behavioural rules such as ‘for coffee, press red button’, while transitioning from ‘knowing what to do’ to ‘actually doing it’. Subtle variation in distributed neural activity patterns reflected rule-specific representations within the ventrolateral prefrontal cortex (VLPFC), confined to instructed stimulus-response learning in contrast to incidental learning involving the same stimuli and responses. VLPFC representations were established right after first-time instruction and remained stable across early implementation trials. More and more fluent application of novel rule representations was channelled through increasing cooperation between VLPFC and anterior striatum. These findings inform representational theories on how the prefrontal cortex supports behavioural flexibility by enabling ad-hoc coding of novel task rules without recourse to familiar sub-routines

## Introduction

The prefrontal cortex (PFC) has been considered crucial for flexibly mastering the abundance of non-routine problems we are often facing ^1-5^. The implementation of completely novel tasks for the very first time is a pivotal example of operating without routine solutions, as those are by definition unavailable ^5,6^. Moreover, it is essential to acquire novel tasks as rapidly as possible to ensure efficient performance and sometimes even physical integrity and ultimately survival. Humans are equipped with the highly developed ability of symbolic communication which is perfectly suited to acquire novel tasks in ‘one shot’ ^7,8^ simply by following explicit instructions. Thereby, more time-consuming and potentially costly trial-and-error learning can be avoided ^9-11^.

Earlier research has generated first insights into the neural basis of instruction-based learning or ‘rapid instructed task learning’ ^12-14^. However, it has remained elusive whether and, if so, how the concrete rules of newly instructed tasks are initially represented in the human PFC right after their first-time instruction. By addressing these questions, the present study set out to inform representational theories of PFC functioning regarding the type and the timescale of task-related information coded within PFC regions. Specifically, we sought to identify distributed neural activity patterns associated with subtle representational differences regarding newly instructed individual rule identities such as ‘if the word BUTTER is displayed on the screen, then flex the middle finger’ or ‘if the word MONKEY is displayed on the screen, then flex the index finger’. To this end, we employed a newly developed multivariate pattern analysis technique (MVPA) specifically calibrated to uncover the rapidly evolving representational dynamics following novel rule instructions ^15^.

Tracking these fine-grained representational dynamics is crucial for a comprehensive understanding of the rapid neural re-organization processes that are taking place right after first-time task instruction. Such rapid neural re-organization processes have been evidenced in terms of both mean activity dynamics ^16-19^ and connectivity dynamics ^20-23^. Specifically, conventional univariate analysis of *mean activity* has shown that lateral PFC engagement was maximal right after instruction, followed by a rapid decline across the first few implementation trials ^16,17,23^. This was paralleled by increasing fronto-striatal *functional connectivity* ^20,24^. Together, these earlier observations suggested that short-term task automatization processes enable increasingly fluent task implementation by support of increasing inter-regional cooperation ^25,26^.

Crucially, however, based on general methodological considerations ^27^, mean activity and connectivity dynamics are uninformative regarding the rule-specific *representational* dynamics being expressed in spatially distributed activity patterns. These can only be uncovered via time-resolved MVPA and by testing a number of alternative scenarios. To start with, newly instructed task rules might be coded within prefrontal cortex right after their first time instruction. If so, the next question then regards the stability of these initially formed representations. One possibility is that initial representations are rapidly fading at the same pace as cognitive control requirements are decreasing (as evidenced by rapidly decreasing mean PFC activity). Alternatively, *stable* prefrontal rule representations might continue to be important for successful task implementation as their more and more fluent application is increasingly channeled through fronto-striatal inter-regional cooperation. A radically different possibility is that novel prefrontal task representations are not yet in place immediately after instruction but are instead being built over the first few implementation trials again based on increasing fronto-striatal cooperation but this time with a leading role of striatal areas. This would be consistent with results in non-human primates during trial-and-error learning showing that successful rule acquisition occurred a few trials *before* rule-specific neural coding could be detected in the lateral PFC ^28,29^.

Importantly, if any of these scenarios could be confirmed empirically, this provided first-time evidence for human PFC representing entirely novel task rules in the initial phase of task practice. This contrasts with existing MVPA studies, which have shown that prefrontal cortex regions flexibly code currently task-relevant information, but this was confined to already *well-familiarized* task features ^4,30,31^. A few pioneering studies have shown that such prefrontal representations are retrieved and re-cycled in the service of newly instructed tasks that rely on, or are recomposed of familiar task elements ^32-34^. Yet, as of now, it remains unknown whether similar regions code entirely novel task rules if these are not composed of familiar task elements.

We conducted two inter-related fMRI experiment both involving a large number of different learning blocks each comprising a new and unique set of instructed stimulus-response (S-R) rules (see Figures 1 and 2). MVPA was used to identify activity patterns sensitive to individual stimulus-response rule identities across the first few implementation trials (see Figure 3). This was done separately for each task block before aggregation across blocks.

**Figure 1.**
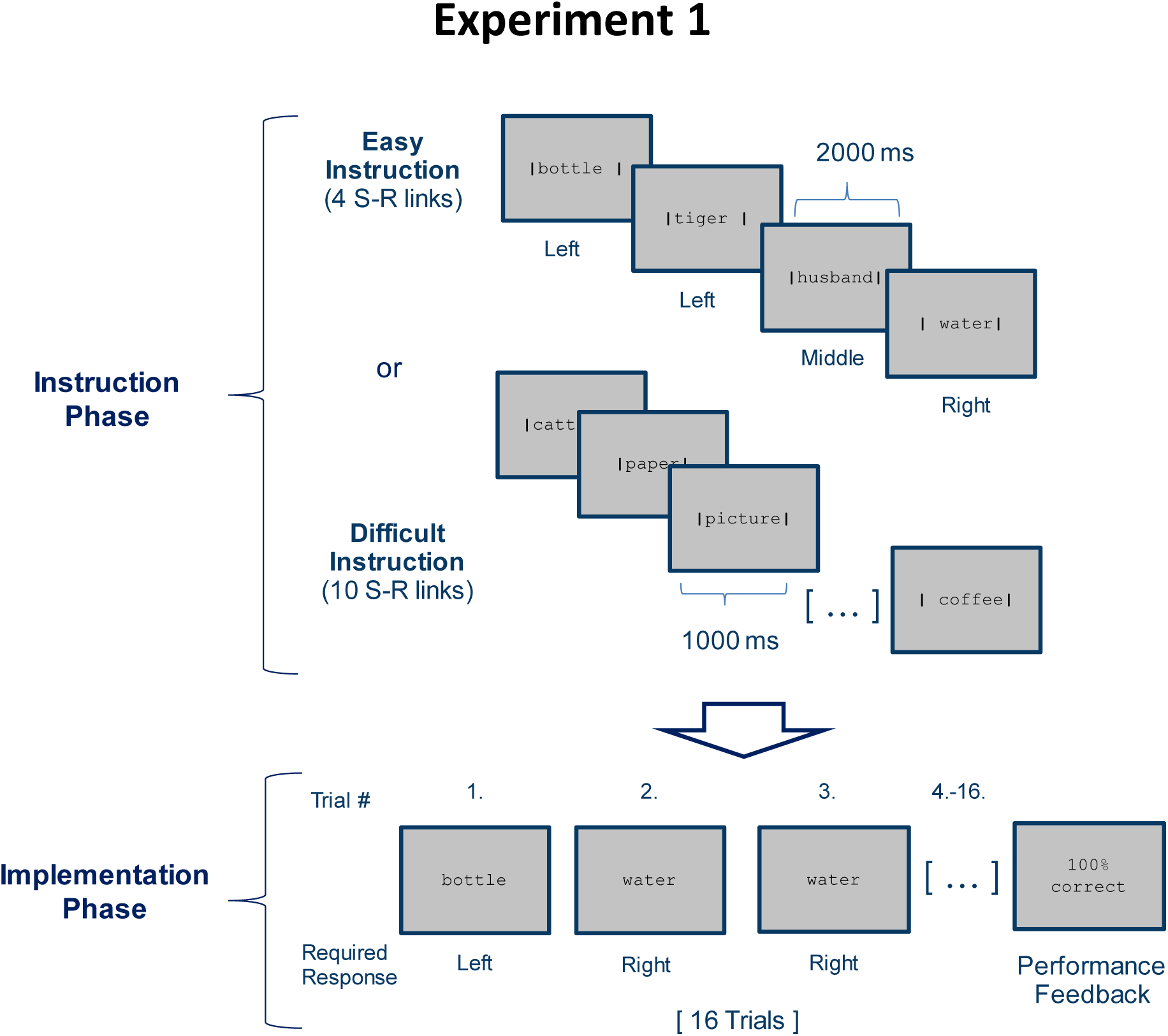
Stimulus-response (S-R) learning task used in experiment 1 exemplarily depicted for one of 18 blocks per condition (easy and difficult). Each block consisted of an instruction phase and an implementation phase. During the instruction phase participants were presented with 4 (easy instruction) or 10 (difficult instruction) pairings between disyllabic nouns and manual responses. The vertical bars framing the nouns indicated the correct response (e.g. Bottle - left). During the subsequent implementation phase (here, selectively shown for the easy condition), each nouns was presented 4 times in random order without the vertical bars and participants had to respond as instructed. Irrespective of S-R rule difficulty (4 vs. 10 nouns in the instruction phase), a constant number of 4 different nouns was presented in the implementation phase. At the end of each block, feedback specifying the percentage of correctly answered trials was displayed.

**Figure 2.**
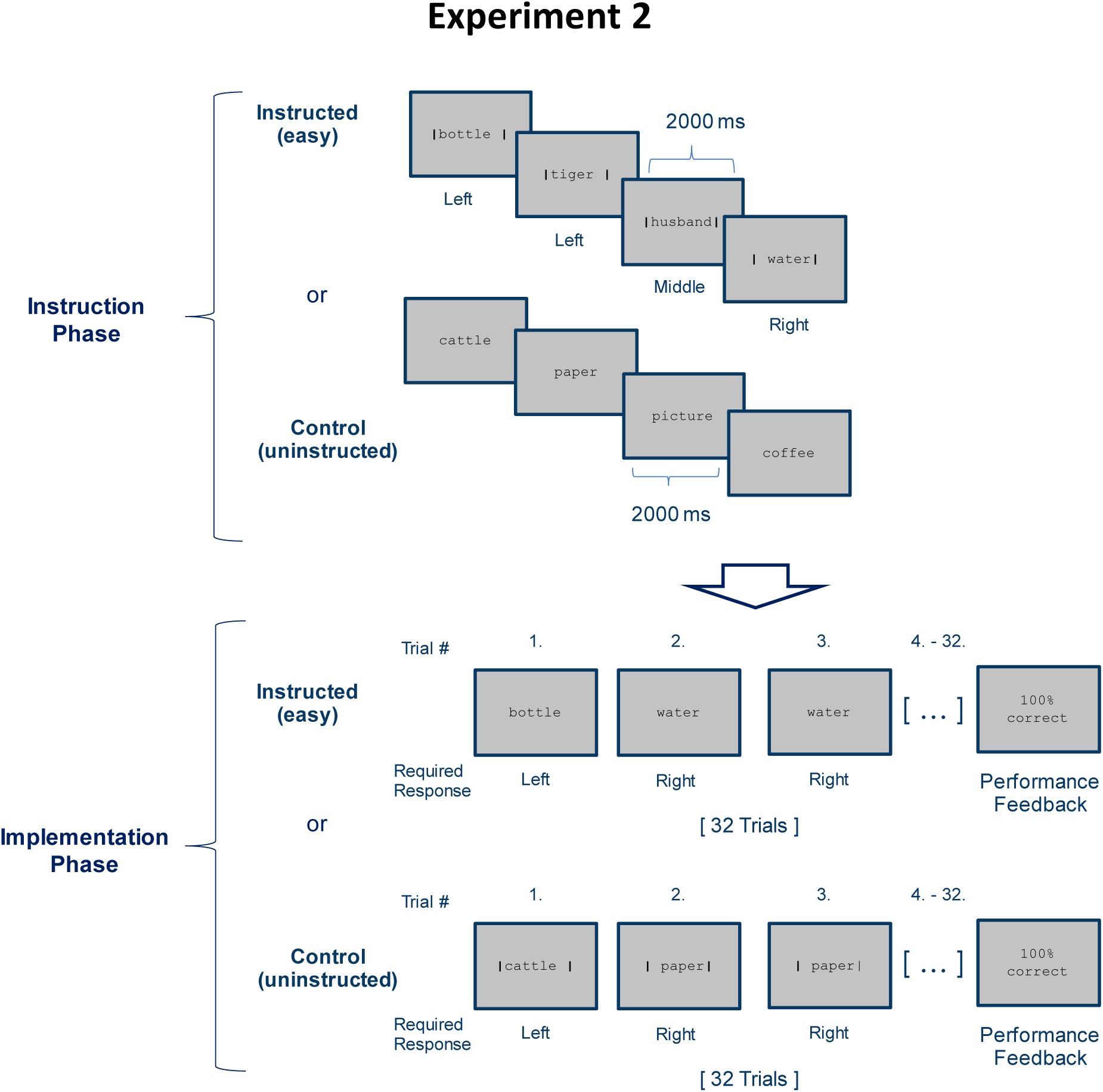
Stimulus-response (S-R) learning task used in Experiment 2 exemplarily depicted for one of 12 blocks per condition (instructed vs. control). As in experiment 1, each block consisted of an instruction phase and an implementation phase. The instructed condition was identical to the easy condition of experiment 1 (i.e., 4 instructed S-R rules) except that each S-R rule needed to be implemented 8 times instead of 4 times. In the control condition the response cues (i.e. the vertical bars) were omitted during the instruction phase and were instead presented during the subsequent implementation phase. At the end of each block, feedback specifying the percentage of correctly answered trials was displayed

**Figure 3.**
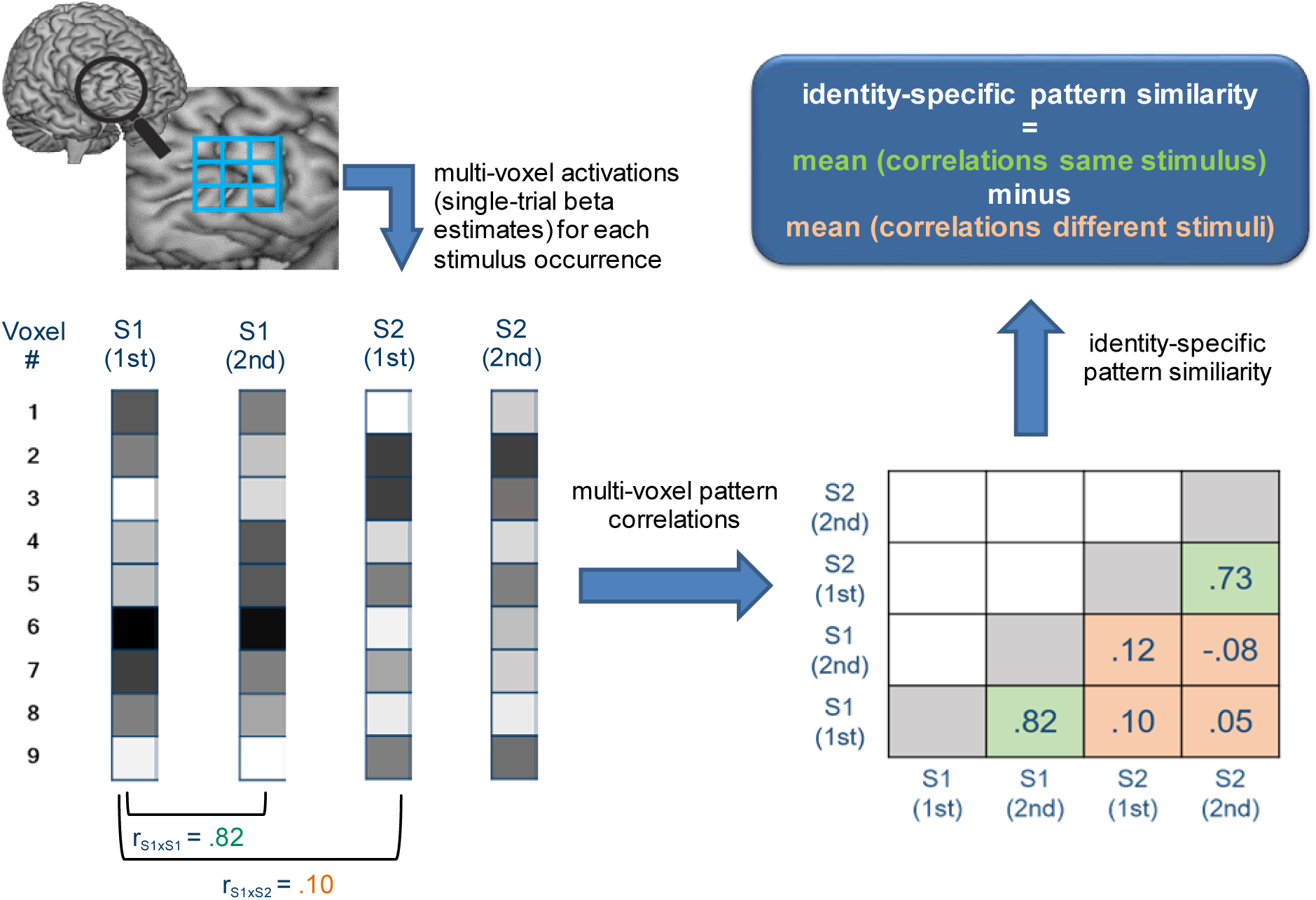
Schematic illustration depicting how identity-specific multi-voxel pattern similarity was computed exemplarily for one learning stage in one learning block. For illustrative purposes, only two stimuli (S1 and S2) each occurring twice are considered here (instead of 4 stimuli in reality). Bottom left: For each stimulus occurrence voxel-wise beta estimates (visualized by grayscale values) are arranged in vectors that constitute the basis of multi-voxel pattern correlations. Bottom right: matrix values depict multi-voxel pattern correlations for all combinations of trials. Green cells denote correlations between same stimuli, orange cells denote correlations between different stimuli. Top right: Identity-specific pattern similarity is defined by significantly greater mean correlations in green cells compared to orange cells.

Besides the primary goal to examine rule-specific representational dynamics, experiment 1 was additionally designed to explore the relationship between the strength or integrity of prefrontal rule representations and the commission of performance errors. To this end, the proportion of performance errors was manipulating by varying the complexity or difficulty of S-R instructions. If performance errors were due to compromised integrity of S-R rule representations, a higher percentage of error trials included in the MVPA following more difficult instructions should imply weaker rule-specific activity patterns ^35,36^. Alternatively, according to the notion of ‘goal neglect’ asserting that ‘knowing’ is not necessarily the same as ‘doing’ ^37,38^, more complex instructions might induce more errors despite largely intact prefrontal rule representations. Instead, more complex instructions might absorb control resources that are then missing to prevent competing (e.g., perseverative) response tendencies from overriding the instructed response. In this case, rule-specific activity patterns should remain unaffected by error rate differences induced by more or less complex instructions.

Experiment 2 was designed as a follow-up to experiment 1 to specifically test the hypothesis that prefrontal cortex representations are confined to *intentional* learning conditions involving instructed stimulus-response rules as compared to an *incidental* learning control condition involving the same contingencies between stimuli and responses but without the necessity to memorize these contingencies for correct performance.

## RESULTS

### Behavioral performance (experiment 1)

RTs and response accuracies from experiment 1 were analyzed with repeated measures ANOVAs. Each ANOVA included the independent variables stimulus repetition (with the levels 1 to 4) and instruction difficulty (with the levels easy and difficult). Greenhouse-Geisser correction was applied where necessary. The results are visualized in Figure 4.

**Figure 4.**
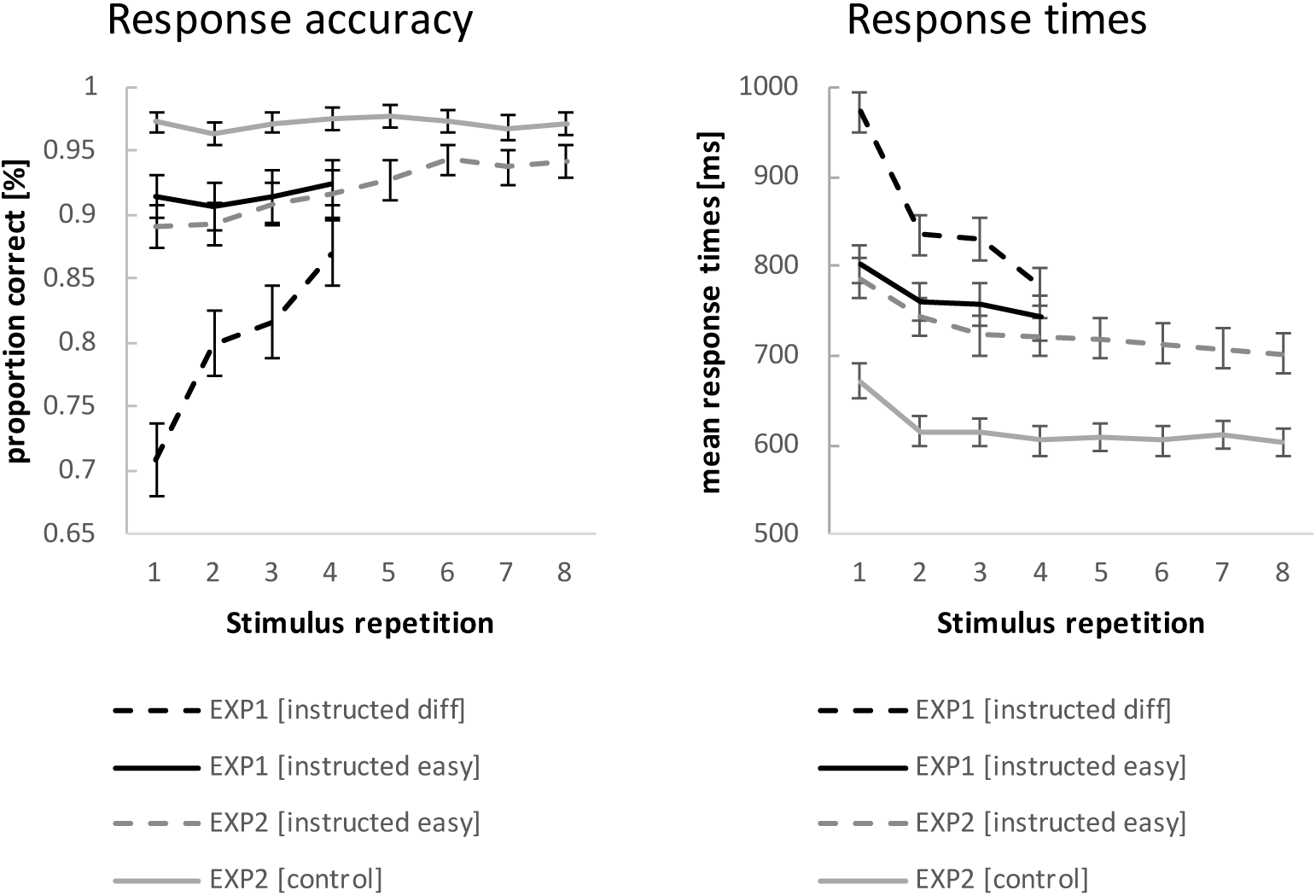
Behavioral performance data for experiment 1 and experiment 2. Error bars represent 90% confidence intervals.

The ANOVA for RTs revealed a significant RT decrease across stimulus repetitions (F_3,192_=224.87; p(F)<.001;*η*_*p*_^*2*^ =.78; linear contrast: F_1,64_=290.67 p(F)<.001; *η*_*p*_^*2*^=.82) which was more pronounced for difficult compared to easy instruction blocks (F_3,192_=137.94; p(F)<.001; *η*_*p*_^*2*^=.68; linear contrast: F_1,64_=252.69; p(F)<.001; *η*_*p*_^*2*^=.80) on top of generally slower RTs (F_1,64_=175.15; p(F)<.001; *η*_*p*_^*2*^=.73). Even at stimulus repetition 4, RTs were still significantly slower for difficult blocks relative to easy blocks (t=4.60; p(t)<.001).

The ANOVA for response accuracies revealed a significant increase in accuracies across stimulus repetitions (F_3,192_=71.43; p(F)<.001; *η*_*p*_^*2*^=.53; linear contrast: F_1,64_=111.52 p(F)<.001; *η*_*p*_^*2*^=.64).This increase was more pronounced for difficult compared to easy instruction blocks (F_3,192_=80.14; p(F)<.001; *η*_*p*_^*2*^=.56; linear contrast: F_1,64_=156.40; p(F)<.001; *η*_*p*_^*2*^=.71) on top of generally higher accuracies for easy blocks than difficult blocks (F_1,64_=202.81; p(F)<.001; *η*_*p*_^*2*^=.76). Even at stimulus repetition 4, accuracies were still significantly higher for easy blocks relative to difficult blocks (t=6.26; p(t)<.001).

Accuracy was positively correlated with the progressive matrices intelligence score, both for easy instructions (r=.32; p=.005 one-tailed) as well as for difficult instructions (r=.35; p=.002 one-tailed). The correlation between the intelligence score and the accuracy difference between easy and difficult instructions showed a trend towards significance (r=-.19; p=.066 one-tailed) indicating that more intelligent participants suffered less from more difficult instructions relative to the easier instructions. Analogous correlations between accuracies and forward and backward simple digit span scores were all non-significant (all p(r)>.14).

### Behavioral performance (experiment 2)

The behavioral data from experiment 2 were analyzed with repeated measures ANOVAs including the independent variables stimulus repetition (with the levels 1 to 8) and instruction type (with the levels instructed and control). Greenhouse-Geisser correction was applied where necessary. The results are visualized in Figure 4. Entering mean response times (RT) as the dependent variable revealed a significant RT decrease across stimulus repetitions (F_7,483_=42.00; p(F)<.001; *η*_*p*_^*2*^=.38; linear contrast: F_1,69_=79.25 p(F)<.001; *η*_*p*_^*2*^=.54), which was more pronounced for instructed blocks compared to control blocks (F_7,483_=4.40; p(F)=.002; *η*_*p*_^*2*^=.06; linear contrast: F_1,69_=6.93; p(F)=.010; *η*_*p*_^*2*^=.09) on top of generally slower RTs (F_1,69_=181.17; p(F)<.001; *η*_*p*_^*2*^=.72). At stimulus repetition 8, RTs were still significantly slower for instructed blocks relative to control blocks (t=12.34; p(t)<.001).

Entering response accuracies as the dependent variable revealed a significant increase in accuracies across stimulus repetitions (F_7,483_=17.51; p(F)<.001; *η*_*p*_^*2*^=.20; linear contrast: F_1,69_=45.88; p(F)<.001; *η*_*p*_^*2*^=.40). This increase was more pronounced for instructed compared to control blocks (F_7,483_=13.95; p(F)<.001; *η*_*p*_^*2*^=.17; linear contrast: F_1,69_=49.10; p(F)<.001; *η*_*p*_^*2*^=.42) on top of generally higher accuracies for control blocks than instructed blocks (F_1,69_=77.18; p(F)<.001; *η*_*p*_^*2*^=.53). At stimulus repetition 8, accuracies were still significantly higher for control blocks relative to instructed blocks (t=5.40; p(t)<.001).

### MVPA (experiment 1)

ROI-based estimates of identity-specific activation patterns from experiment 1 were submitted to a 4-factorial repeated-measures ANOVA including the independent variables learning stage (early vs. late) and difficulty (easy vs. diff) and additionally region (VLPFC vs. DLPFC) and hemisphere (left vs. right) in order to adequately account for potential regional differences ^39^. The results are visualized in Figure 5. This analysis yielded a significant overall identity-specific pattern similarity effect (F_1,64_=7.9; p(F)=.006; *η*_*p*_^2^ = .11) which was significantly stronger for VLPFC than DLPFC (F_1,64_=13.1; p(F)<.001; *η*_*p*_^2^ = .17).

**Figure 5.**
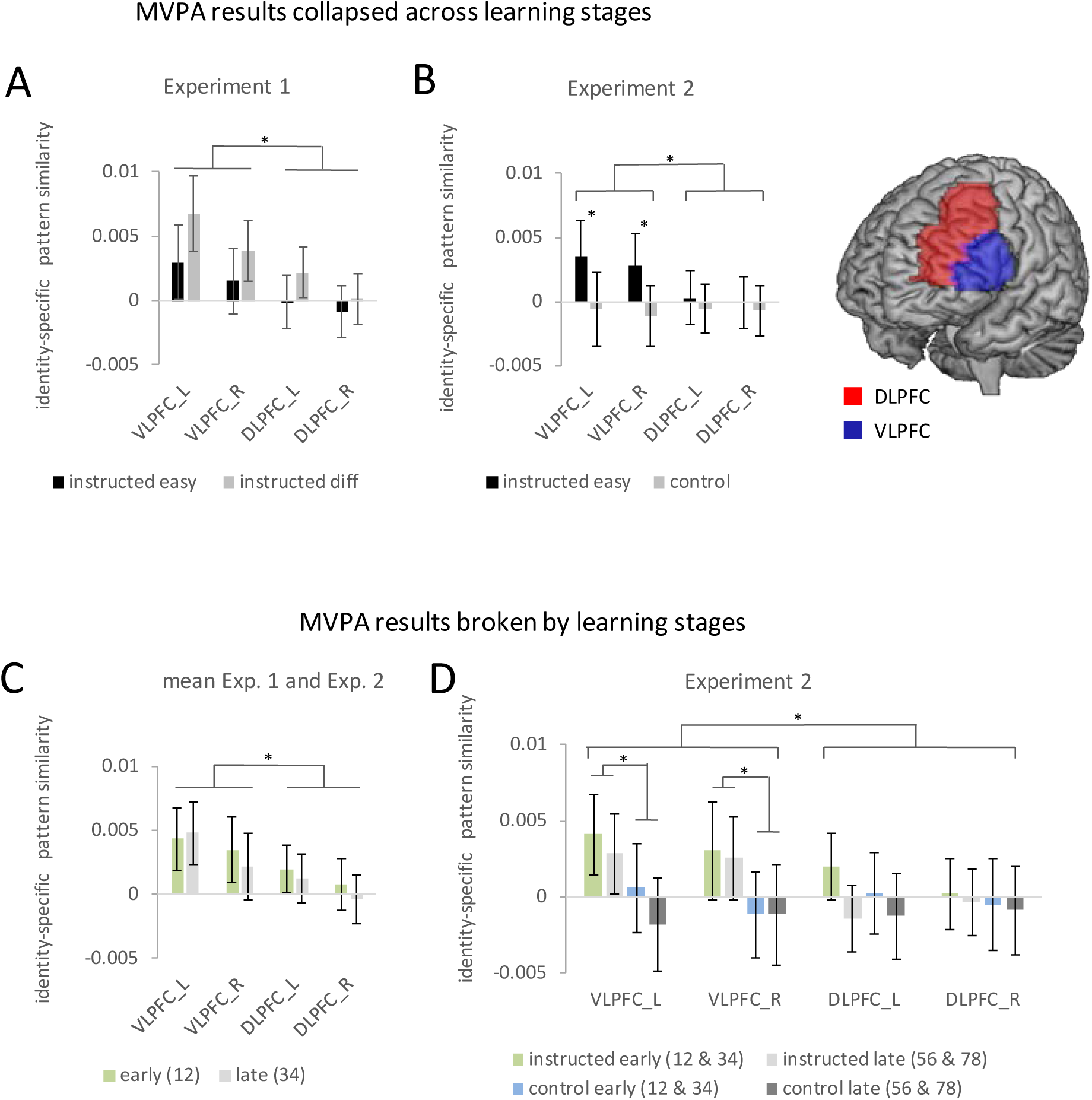
Summary of the ROI-based MVPA results for experiments 1 and 2. Error bars represent 90% confidence intervals. Significant differences are indicated by asterisks. (A) Identity-specific pattern similarities in experiment 1 collapsed across learning stages. (B) Identity-specific pattern similarities in experiment 2 collapsed across learning stages. (C) Identity-specific pattern similarities collapsed across experiments 1 and 2 broken by learning stages. Early learning stage pattern similarities are based on stimulus repetitions 1 and 2 whereas late learning stage pattern similarities are based on stimulus repetitions 3 and 4. (D) Identity-specific pattern similarities for experiment 2 broken by learning stages. Early learning stage pattern similarities are based on aggregated values for stimulus repetitions 1 and 2 and stimulus repetitions 3 and 4. Late learning stage pattern similarities are based on aggregated values for stimulus repetitions 3 and 4 and stimulus repetitions 7 and 8.

There were no significant effects involving learning stage but a trend towards a smaller effect for late vs. early in the VLPFC compared to the DLPFC (F_1,64_=3.6; p(F)=.064; *η*_*p*_^*2*^=.053). In order to test whether this trend might point towards a ‘true’ but small effect that was missed due to insufficient statistical power, we conducted an additional more powerful analysis by collapsing data across experiments 1 and 2. However, this analysis again did not produce reliable evidence for a significant influence of learning stage (for details see further below). Also, there were no significant effects involving difficulty. If anything, contrary to the prediction of weakened rule representations, there was a trend towards a *stronger* identity-specific pattern similarity effect in the difficult condition compared to the easy condition (F_1,64_=3.3; p(F)=.074; *η*_*p*_^*2*^=.049).

The ROI-based findings were confirmed by searchlight-based MVPAs within each ROI, revealing a significant overall identity-specific pattern similarity effect specifically within the left VLPFC (MNI: −48 5 23; t=5.09; p_FWE_<.001 and MNI: −45 32 11; t=4.66; p_FWE_<.001) and a trend in the same direction within the right VLPFC (MNI: 48 8 11; t=3.16; p_FWE_<.074). Again, there were no significant effects involving learning stage or difficulty.

On the whole brain level, the searchlight MVPA confirmed for the left VLPFC that the overall identity-specific pattern similarity effect was significant even after correction for the whole-brain volume (MNI: −48 5 23; t=5.09; p_FWE_=.005 and MNI: −45 32 11; t=4.66; p_FWE_=.028). Additionally, this analysis revealed significant whole-brain-corrected effects in the left sensorimotor cortex (MNI: −39 −25 53; t=9.28; p_FWE_<.001) and in the left visual cortex (MNI: −15 −91 −7; t=6.52; p_FWE_<.001). There were no significant effects involving learning stage or difficulty. These findings are as expected and consistent with the coding of stimulus identity in the visual cortex and response identity in the left sensorimotor cortex, respectively (see Figure 6).

**Figure 6.**
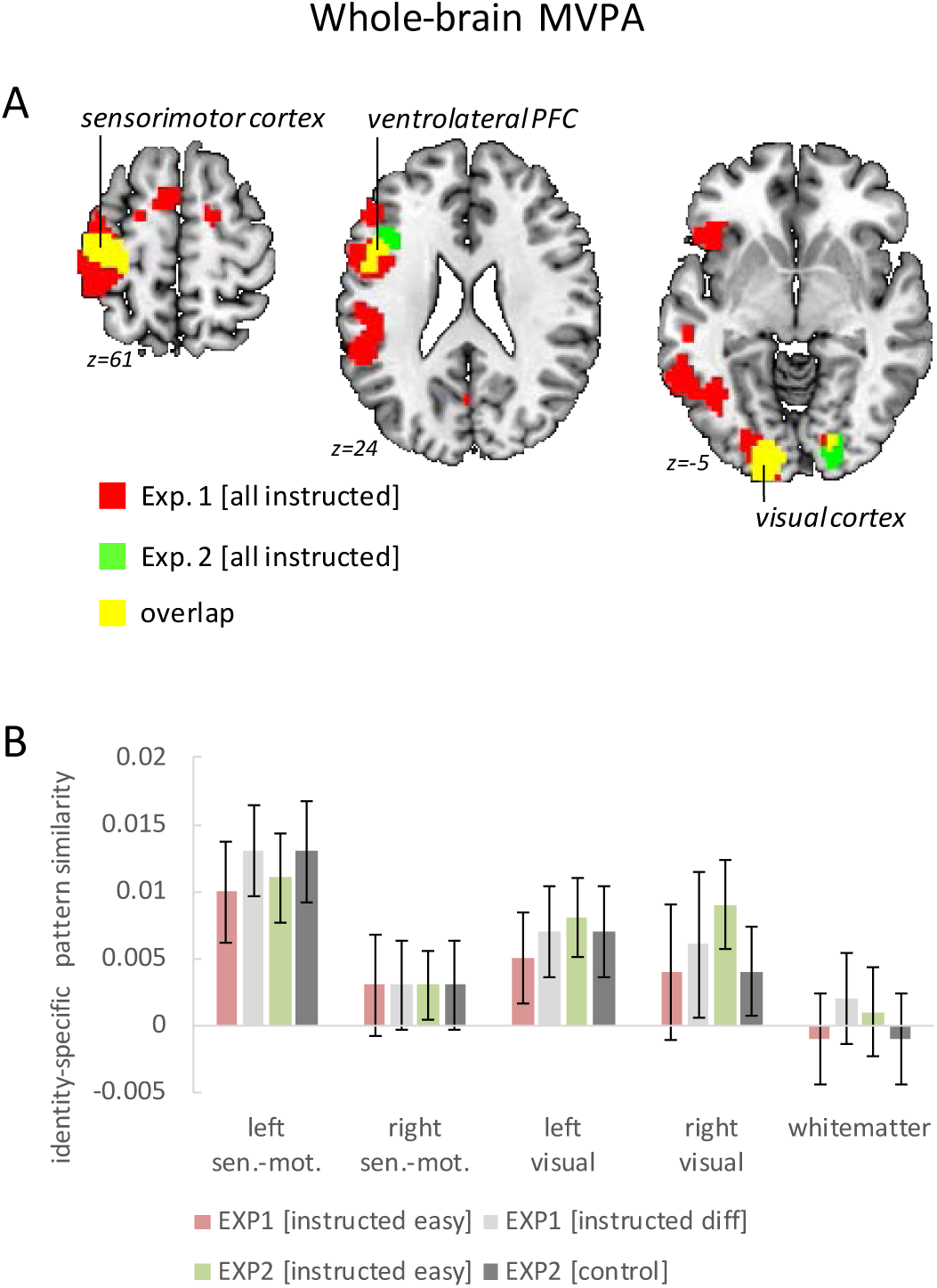
Results of the whole-brain searchlight MVPA testing for overall identity-specific pattern similarity effects. (A) Horizontal brain slices depicting the findings for the left sensorimotor cortex, the ventro-lateral PFC, and the visual cortex. For display purposes the map shows voxels with p<.001 uncorrected. (B) Pattern-similarity effects broken by instruction difficulty (exp. 1) or instruction type (exp. 2). In addition to sensorimotor cortices and visual cortices, the white-matter volume is included as a control region to highlight the absence of analysis bias. For a comprehensive summary of ventro-lateral PFC results see Fig. 5. Error bars represent 90% confidence intervals.

### MVPA (experiment 2)

Instead of reflecting S-R rule-specific representations, the findings of experiment 1 could in principle reflect representations of stimulus identity or response identity alone. In fact, this was very likely the case for the left sensorimotor cortex (response identity) and for the visual cortex (stimulus identity). Moreover, it remains unclear whether MVPA effects observed in higher-order brain regions like the VLPFC really indicated *intentionally* encoded S-R rules according to the explicit instruction or rather *incidentally* encoded S-R rules. To clarify this, experiment 2 included a control condition which was identical in terms of information content regarding stimuli, responses, and S-R contingencies. The only difference was that novel S-R rules were not required to be actively retrieved and hence intentional encoding was discouraged (while incidental encoding was still possible during the implementation phase).

As in experiment 1, ROI-based estimates of identity-specific activation patterns were submitted to a 4-factorial repeated-measures ANOVA including the independent variables learning stage (early vs. late), instruction type (instructed vs. control), and additionally region (VLPFC vs. DLPFC) and hemisphere (left vs. right) in order to adequately account for potential regional differences ^39^. The results are visualized in Figure 5. Note that different from experiment 1, this time the early learning stage comprised the mean across identity-specific pattern similarities computed for stimulus repetitions 1 and 2 and stimulus repetitions 3 and 4, respectively. The late learning stage comprised the mean across identity-specific pattern similarities computed for stimulus repetitions 5 and 6 and stimulus repetitions 7 and 8, respectively. The ANOVA yielded a significant main effect of instruction type (F_1,69_=4.49; p(F)=.038; *η*_*p*_^2^ = .061) which was significantly stronger for VLPFC than DLPFC (F_1,69_=6.91; p(F)<.011; *η*_*p*_^2^ = .091). There was no significant effect involving learning stage. Unlike experiment 1, there was not even a trend towards an influence of learning stage when testing the relevant interaction involving stage, region, and instruction type (F_1,69_=.22; p(F)=.641; *η*_*p*_^2^ = .003). Note that similar results were obtained when learning stage comprised all four non-aggregated levels (i.e. based on stimulus repetitions 1/2, 3/4, 5/6, and 7/8) instead of the two aggregated levels used in the primary analysis.

Thus, the ROI-based MVPA confirmed the findings from experiment 1 and importantly showed that identity-specific MPVA effects are indeed specific of instructed S-R rules as compared to the control condition involving the same stimuli and responses under incidental S-R learning conditions. Notably, again consistent with experiment 1, the MVPA effects for instructed S-R learning relative to control were significantly stronger in the VLPFC compared to the DLPFC where an effect was virtually absent.

These ROI-based findings were confirmed by searchlight-based MVPAs within each ROI revealing stronger identity-specific pattern similarity effects in the instructed condition than in the control condition specifically within the left VLPFC ROI (MNI: −36 17 26; t=3.51; p_FWE_=.025) and a trend in the same direction also within the right VLPFC (MNI: 60 14 14; t=2.90; p_FWE_=.134). On the whole brain level, the searchlight MVPA did not reveal additional regions exhibiting a main effect of instruction type. Neither learning stage nor instruction-type had a significant influence on the searchlight results. Notably, testing for identity-specific activation patterns collapsed across instructed blocks and control blocks revealed the expected effects for both conditions alike in the sensorimotor cortex (MNI: −39 −25 50; t=10.48; p_FWE_<.0001) and the visual cortex (MNI: −15 −91 −4; t=5.1; p_FWE_=.003 and MNI: 21 −88 −4; t=4.67; p_FWE_=.02). These results are depicted in Figure 6 and confirm the findings from experiment 1. Importantly, different from the VLPFC findings which were highly specific for the instructed learning condition, visual cortex and sensorimotor cortex exhibited – as expected – comparable effects both in the instructed learning condition as well as in the control condition. This is consistent with representations of stimulus identity and response identity, respectively.

### MVPA (collapsed across experiments 1 and 2)

Experiment 1 exhibited a non-significant trend towards weaker identity-specific pattern similarity for the late learning stage relative to the early learning stage. In order to test whether this trend might point towards a ‘true’ but small effect that was missed due to insufficient statistical power, we conducted an additional more powerful analysis based on data from both experiments. Data from experiments 1 and 2 were jointly analyzed including all the instructed conditions (i.e., omitting the control condition from experiment 2) for the early stage spanning stimulus repetitions 1 and 2 and the late stage spanning stimulus repetitions 3 and 4 (i.e., omitting stimulus repetitions 5/6 and 7/8 from experiment 2). The results are visualized in Figure 5C.

ROI-based estimates of identity-specific pattern similarity were submitted to a 3-factorial repeated-measures ANOVA including the independent variables learning stage (early vs. late), region (VLPFC vs. DLPFC), and hemisphere (left vs. right). Not surprisingly, this ANOVA again yielded a significant overall identity-specific pattern similarity effect (F_1,134_=9.37; p(F)=.003; *η*_*p*_^*2*^ = .065) which was significantly stronger for the VLPFC than the DLPFC (F_1,134_=14.04; p(F)<.001; *η*_*p*_^*2*^ = .095). Most importantly, refuting the preliminary trend observed in experiment 1, this latter effect was *not* significantly affected by learning stage (F_1,134_=.47; p(F)=.49; *η*_*p*_^*2*^ = .004). All other ANOVA effects involving learning stage were also non-significant (all p>.40). Hence, overall, it seems relatively safe to conclude that identity-specific pattern similarity effects in the VLPFC are stable across learning stages.

### Univariate analysis (experiment 1)

A complementary ROI-based univariate analysis for experiment 1 was based on condition-specific mean activity estimates which were submitted to a 4-factorial repeated-measures ANOVA including the independent variables stimulus repetition (1 to 4), difficulty (easy vs. diff), region (VLPFC vs. DLPFC), and hemisphere (left vs. right). The results are visualized in Figure 7. There was a significant main effect of stimulus repetition (F_3,192_=62.38; p(F)<.001; *η*_*p*_^*2*^ = .29) reflecting a general linear activation decrease (linear contrast: F_1,64_=187.11; p(F)<.001; *η*_*p*_^*2*^ = .39). A significant three-way interaction involving difficulty, region, and hemisphere (F_1,64_=7.20; p(F)=.009; *η*_*p*_^*2*^ = .10) reflected stronger activation in the difficult condition relative to the easy condition which was especially pronounced in the left DLPFC. This was further qualified by a significant four-way interaction additionally including stimulus repetition (F_3,192_=5.96; p(F)=.001; *η*_*p*_^*2*^ = .09) reflecting a linearly decreasing influence of difficulty which was especially pronounced in the left VLPFC (linear contrast: F_1,64_=9.82; p(F)=.003; *η*_*p*_^*2*^ = .13).

**Figure 7.**
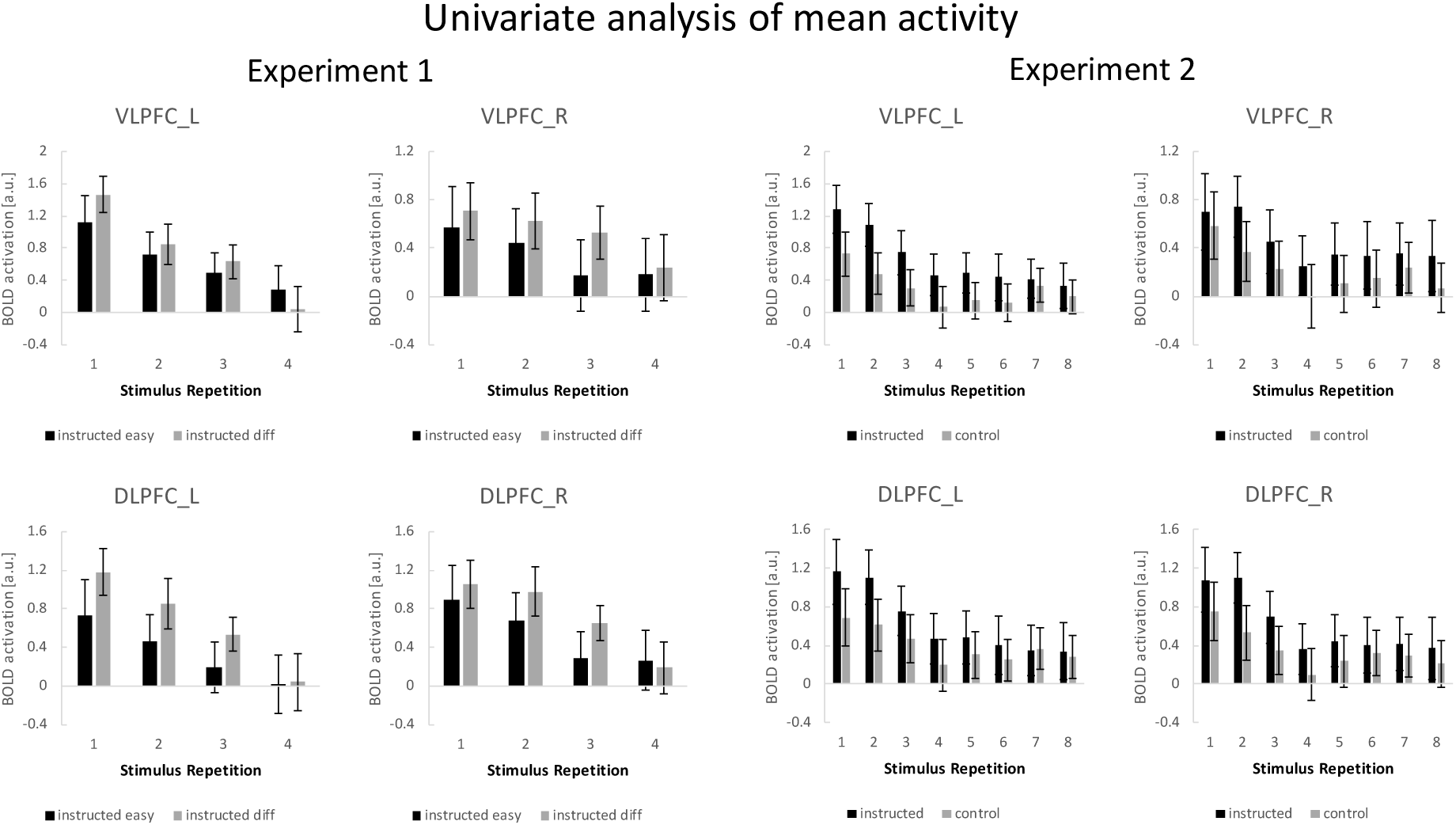
Summary of the ROI-based mean activity results for experiments 1 and 2. Error bars represent 90% confidence intervals.

### Univariate analysis (experiment 2)

A complementary ROI-based univariate analysis for experiment 2 was based on condition-specific mean activity estimates which were submitted to a 4-factorial repeated-measures ANOVA including the independent variables stimulus repetition (1 to 8), instruction type (instructed vs. control), region (VLPFC vs. DLPFC), and hemisphere (left vs. right). The results are visualized in Figure 7. This ANOVA yielded significant main effects of instruction type (F_1,69_=5.31; p(F)=.024; *η*_*p*_^*2*^ = .071) and stimulus repetition (F_7,483_=14.51; p(F)<.001; *η*_*p*_^*2*^ = .174) indicating generally higher activation for the instructed blocks relative to the control blocks and generally decreasing activation across stimulus repetitions. Notably, however, a significant four-way interaction between all independent variables (F_7,483_=4.27; p(F)=.002; *η*_*p2*_ = .058) indicated that the stronger activation for instructed blocks was linearly decreasing across stimulus repetitions, but to a different extent across ROIs and particularly pronounced for the left VLPFC (linear contrast: F_1,69_=10.54; p(F)=.002; *η*_*p*_^*2*^ = .133).

### Functional connectivity analysis (experiment 2)

Previous studies have reported *increasing* connectivity across stimulus repetitions between the LPFC and the anterior striatum under instruction-based learning conditions ^20,24^. The study design of the present experiment 2 offers the unique opportunity to explicitly test whether this effect is specific of instructed learning blocks compared to control blocks. Such a finding would additionally inform the MVPA results by suggesting that the repeated application of newly established VLPFC rule representations might be increasingly channelled through inter-regional cooperation between the VLPFC and the anterior striatum.

Analogously to the earlier studies, we tested for a stronger functional connectivity increase from early learning trials (stimulus repetitions 1 and 2) to late learning trials (stimulus repetitions 7 and 8). The results are visualized in Figure 8. Using the left VLPFC as seed region, we specifically tested for significant beta-series correlation effects within an anatomically defined basal ganglia ROI comprising all of caudate nucleus, putamen, and pallidum. This revealed the predicted effect in the anterior striatum (MNI: −6 14 −4; t=4.08; p(t)=.018 and MNI: 6 14 −7; t=4.11; p(t)=.016; FWE-corrected for the basal ganglia volume). There were no additional regions identified after correction for the whole brain volume. Note that also the striatal activation dynamics were as expected based on previous studies. Specifically, as visualized in Figure 8, there was a significant linear activation increase for instructed learning blocks relative to the control blocks for the anterior striatum cluster identified in the connectivity analysis (MNI: 6 14 −7; t=4.35; p(t)=.001 and MNI: −9 20 −7; t=4.53; p(t)=0.001 and MNI: 18 23 −7; t=4.96; p(t)<.001; all FWE-corrected for the ant. striatum volume). Together, these findings lend further support for an early practice-related increase in anterior striatal activity and connectivity specifically under instruction-based learning conditions as has been debated recently ^23,26^.

**Figure 8.**
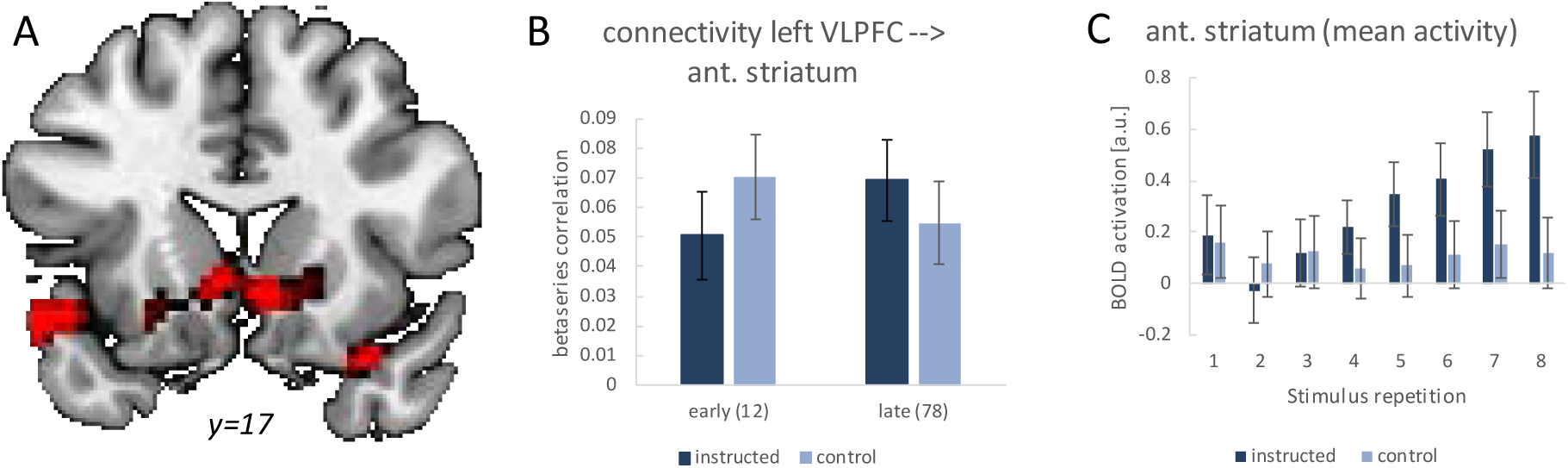
Summary of the functional connectivity analysis results for the left VLPFC seed region based on single-trial beta-series correlations. The analysis tested for a functional connectivity increase from early learning trials (stimulus repetitions 1 and 2) to late learning trials (stimulus repetitions 7 and 8) which was stronger for instructed learning blocks than control blocks. (A) Visualization of the significant effect in the anterior striatum. For display purposes the map shows voxels with p<.001 uncorrected. (B) The detailed connectivity pattern for the anterior striatum cluster. (C) Mean activations at each stimulus repetition level based on a conventional univariate analysis for the anterior striatum cluster. Error bars represent 90% confidence intervals.

## DISCUSSION

The key finding from the time-resolved MVPA is that rule identity-specific representations were established immediately after the first-time instruction of completely novel S-R rules and continued to be involved throughout the first few implementation trials. This effect was highly specific for the VLPFC and it was clearly not detectable in the DLPFC. Importantly, we could show that identity-specific pattern similarity effects in the VLPFC were indeed confined to the intentional learning condition involving instructed S-R rules as compared to a control condition involving the same stimuli and responses under incidental S-R learning conditions. Moreover, the stable representation of rules within the VLPFC was paralleled by an increasing functional coupling between the VLPFC and the anterior striatum. This seems to suggest that the more and more fluent application of newly established rule representations is increasingly channelled through inter-regional cooperation during an early phase of short-term task automatization ^25^. Interestingly, the stability of rule representations within the VLPFC stands in stark contrast to the rapidly decreasing mean activity revealed by the univariate analysis. Moreover, while the multivariate pattern similarity effect was tightly confined to the VLPFC, the decreasing mean activity spread across VLPFC and DLPFC. This re-emphasizes the insight that mean activity results are unsuited to draw meaningful conclusions regarding the representational content of brain regions ^27^ and sets into perspective somewhat over-interpreted univariate analysis results we have reported earlier ^16^.

The stability of VLPFC rule representations in the present study is distinctly different from observations reported by earlier electrophysiological studies in non-human primates in the context of trial-and-error learning ^28,29^. Those studies found that successful rule acquisition occurred (often quite abruptly) a few trials *before* rule-specific neural coding could be detected in the LPFC. In other words, even though overt behavior clearly suggested that a novel S-R rule had been successfully acquired, the lateral PFC did not seem to initially represent this rule. By contrast, anterior caudate neurons directly reflected improvements in behavioral accuracy ^28^. This suggests that under trial-and- error learning conditions and in non-human primates the anterior caudate rather than the lateral PFC might be the place where novel task rules are initially represented. Further research is necessary to clarify whether these differences in representational dynamics are due to (i) differences between trial-and-error learning and instruction-based learning, (ii) general differences between species, or (iii) regional differences between the VLPFC region (area BA 44/45) identified in the present study and the DLPFC region (area BA 9/46) selectively examined in the electrophysiological studies.

### Representations of newly instructed rules and familiar rules

Our primary aim to track the initial representational dynamics of newly instructed task rules naturally required an ‘aggregation-free’ MVPA approach based on single-trial estimates associated with the trial-by-trial coding of individual S-R rule identities. This contrasts with earlier MVPA studies which relied in one way or the other on aggregation schemes either across time ^40-45^ or across individual rule identities ^32,34,35,46,47^. Aggregation across individual rule identities improves signal-to-noise ratio regarding representations of task features on a more abstract level, but this generalization comes at the cost of losing specificity regarding individual rule identities. Similarly, aggregation across time, which typically involves aggregation across a large number of trials per rule identity, improves signal-to-noise ratio regarding each rule identity. Hence, this approach is obviously unsuited to track rapidly evolving representational dynamics spanning only a few trials, but is instead suited to examine representations involving well-familiarized task rules or to track slow learning processes evolving across blocks of large numbers of trials per rule identity.

Such aggregation-based studies could demonstrate that information regarding well-familiarized rule identities is flexibly represented within the prefrontal cortex under conditions that often require the prioritized implementation of one currently relevant task over competing alternative tasks ^33,35,40-43,48,49^. This is consistent with similar findings reported in electrophysiological studies in non-human primates ^3,50-52^. Overall these studies nicely show that the prefrontal cortex flexibly codes anything of current task relevance, including information regarding task-relevant stimuli, responses, perceptual and conceptual categories, and transformation rules like those required in typical stimulus-response tasks (Crittenden, Mitchell, & Duncan, 2016; Duncan, 2010; Fedorenko et al., 2013; Woolgar et al., 2016). However, unlike the present study, these earlier conclusions were restricted to already well-familiarized task features, and could hence not tell whether prefrontal representational flexibility also extends to completely novel tasks.

A number of pioneering MVPA studies specifically focusing on instruction-based learning could show that representations of *familiar* task features can be retrieved and re-cycled in the service of newly instructed tasks. One such study by Muhle-Karbe et al. ^34^ identified LPFC activity patterns associated with highly familiar categorization routines regarding house pictures vs. face pictures – but, importantly, not regarding the concrete stimulus-response rules (e.g., the instructed responses for each of two different faces) underlying a multitude of individual face or house categorization tasks each involving a unique set of stimuli. Similar conclusions apply to a related study ^53^. Another approach pursued by Cole et al. ^32,33^ provided evidence for the principle of ‘rule compositionality’ see also ^54,55^. They showed that distributed activity and connectivity patterns of familiar task elements (e.g. same/different judgement or semantic categorization) were re-activated when these task elements were later combined with each other to construct a multitude of novel tasks defined by their specific combination and applied to a set of novel stimuli. Importantly, MVPA was based on aggregation over all those novel task compositions that shared one specific rule element. Hence, while being highly informative regarding the question of rule compositionality, this type of study does not speak to the question of how the *identities* of individual novel task compositions might be represented in the brain. The same holds for identities of entirely novel tasks that are not composed of familiar task elements and this is exactly the question that was answered by the present study.

### Rule representations and the complexity of S-R instructions

An additional goal of experiment 1 was to explore the relationship between the strength or integrity of prefrontal rule representations and the extent of performance errors as a function of the complexity of S-R instructions. One of our original hypotheses was inspired by previous study results 35,36 and presumed that most errors would be committed due to damaged representations of the originally instructed S-R rules. Hence, a higher proportion of errors in the more difficult condition should be associated with a weaker identity-specific pattern similarity effect. However, if anything there was a non-significant trend towards a stronger identity-specific pattern similarity effect in the more difficult condition. A possible explanation of this null finding is based on a radically different account related to the notion of goal-neglect ^37,56^ and could explain why the strength or integrity of prefrontal cortex representations remained unaffected by differences in instructed rule complexity. Alluding to the difference between ‘knowing’ and ‘doing’ ^37,57^, more complex instructions might induce more errors despite largely intact VLPFC representations. Instead, error rate might increase due to failures to correctly implement (‘doing’) correctly retrieved rules (‘knowing’). This is consistent with VLPFC housing ‘declarative’ rather than ‘procedural’ rule representations ^58^ possibly related to the concept of an ‘episodic buffer’ within working memory ^37,59^. Implementation errors despite ‘knowing better’ might occur when more complex instructions absorb additional control resources that are then lacking in order to prevent competing (e.g., perseverative) response tendencies from overriding the instructed correct response. Such a resource ‘depletion’ account would predict generally increased control effort following more complex instructions – including correctly performed trials. This prediction is indeed supported by the univariate analysis which revealed stronger mean activity in prefrontal cortex for more complex instruction blocks (paralleled by significantly increased response times). Additional support comes from the finding that response accuracies were positively associated with Raven’s progressive matrices intelligence scores but not with simple working memory span. This seems to suggest that response errors were not so much related to the inability to memorize the instructions but rather to a more general cognitive control deficit reflected by the intelligence score. This is consistent with the observation that general intelligence is associated with goal neglect ^37^.

## General conclusions

Our findings are suited to inform representational theories on how the prefrontal cortex supports behavioural flexibility. Specifically, we demonstrated that the VLPFC achieves flexibility not only by recycling established sub-routines in the service of novel task requirements but also by enabling the ad-hoc coding of novel task rules instantaneously after their first-time instruction and without recourse to established sub-routines. This refutes alternative accounts that would have predicted an incremental process of rule formation in the prefrontal cortex possibly driven by leading signals generated by striatal areas. On the contrary, our findings suggest the reverse relationship between VLPFC and anterior striatum where the application of instantaneously established prefrontal rule representations is increasingly channelled through inter-regional cooperation with the anterior striatum. Future research is however needed to further clarify the relationship between striatal areas and prefrontal areas with respect to novel task learning under a greater variety of circumstances. In particular, this might include systematic explorations regarding (i) different types of intentional learning such as trial-and-error learning vs. instruction-based learning, (ii) different age groups or different species, and (iii) different time scales.

## Methods

### Participants

The sample for experiment 1 consisted of 65 human participants (32 female, 33 male; mean age: 24.2 years, range 19-33 years). Three additional subjects could not be used due to incomplete data collection. Part of the present dataset was used in a previous methods-oriented paper ^15^. The sample for experiment 2 consisted of 70 human participants (39 female, 31 male; mean age: 23.9 years, range 19-33 years). Two additional subjects could not be used due to incomplete data collection. All participants were right-handed, neurologically healthy and had normal or corrected vision. The experimental protocol was approved by the Ethics Committee of the Technische Universität Dresden and conformed to the World Medical Association’s Declaration of Helsinki. All participants gave written informed consent before taking part in the experiment and were paid 10 Euros per hour for their participation or received course credit.

### Tasks

Both experiments were based on modified versions of an established instruction-based learning paradigm ^16^. Generally, the participants worked through a series of different novel tasks blocks. In each task block they were required to memorize novel task instructions during an initial *instruction phase* during which response execution was not yet required. The instruction phase was followed by a manual *implementation phase* requiring task execution on a trial-by-trial basis by retrieving the previously encoded task rules from memory. In both experiments a task instruction comprised a set of novel stimulus-response (S-R) rule identities. The term ‘rule identity’ refers to a specific link between one unique stimulus and the response assigned to that stimulus. Each set of stimuli comprised either 4 or 10 written disyllabic German nouns which were mapped onto either 2 or 3 different manual button press responses (index, middle, or ring finger of the right hand). The number of responses was varied in order to encourage the memorization of all S-R rules and to avoid excessive use of short-cuts like ‘these two stimuli require response A, hence all other stimuli require the other response’ ^60^. The number of task blocks requiring either 2 or 3 different responses was equally distributed across the different instruction conditions (easy/difficult in experiment 1 and instructed/control in experiment 2).

The start of an impending instruction phase was announced by the German word for ‘memorize’ (‘Einprägen’) displayed in red for 2 s, followed by the presentation of the first instructed noun. The start of the instruction phase announcement was delayed by a variable delay of 2 or 4 s relative to the start of a new measurement run or relative to the end of the preceding implementation phase. During instruction, the novel nouns were presented in rapid succession framed by two vertical bars to the left and to the right of the noun (see Figure 1). If a noun was closer to the left vertical bar, this indicated an index finger response. If a noun was closer to the right vertical bar, this indicated a ring finger response. If a noun was equally close to both vertical bars, this indicated a middle finger response. We only recruited right-handed subjects who were asked to use the right hand fingers for responding.

During the manual implementation phase which directly followed the instruction phase, the stimuli were presented in pseudo-random order such that each stimulus was presented 4 times (experiment 1) or 8 times (experiment 2). Each implementation phase was announced by the German word for ‘implement’ (‘Ausführen’) displayed in green for 2 s. There was no performance feedback after individual trials to avoid interference with reinforcement learning. The SOA varied randomly between 2 and 4 s in 0.5 s steps. The SOA interval was inserted *before* the start of a new trial to ensure that there was also random jitter between the end of the instruction phase and the beginning of the first implementation trial. After a variable delay of 2 or 4 s relative to the end of the last trial, the implementation phase ended with a display (2 s) of the mean performance accuracy computed across the preceding trials.

### Experiment 1 specifics

The aim of experiment 1 was twofold. First, we wanted to identify rule-specific neural representations with maximal statistical power and focused on the earliest phase of learning. We therefore realized a large number of 36 unique learning blocks each comprising only 4 repetitions of each of four stimuli. Second, we wanted to explore the relationship between the strength or integrity of prefrontal representations and the commission of performance errors. We therefore manipulated the complexity or difficulty of S-R instructions. The two difficulty conditions only differed regarding the number of instructed S-R rules (4 vs. 10) but not regarding the number of actually implemented S-R rules (always 4). In the difficult condition, 10 nouns were instructed and each was displayed for 1 s. In the easy condition, 4 nouns were instructed and each was displayed for 2 s. With respect to the subsequent implementation phase, the two conditions were identical, i.e., in either case, 4 nouns were presented. The subset of 4 out of 10 instructed nouns presented during the implementation phase of the difficult condition was selected such that 2 or 3 different responses were required equally often. Participants performed 18 blocks of each condition in pseudo-randomized order, which took approximately 40 minutes. Measurements were taken in three consecutive runs of ca. 13 min duration, each comprising 6 blocks of each difficulty condition. Also, the random delay before the start of each novel instruction phase and the delay before performance feedback was pseudo-randomized such that each SOA level occurred equally often for each difficulty condition. Before entering the scanner each participant completed a short practice session comprised of one novel task block for each difficulty condition with a separate stimulus set not used during the main experiment.

After completion of the instruction-based learning experiment in the scanner, participants performed a computerized simple digit span task to determine individual simple working memory span scores ^61^. This score was chosen to obtain a relative pure measure of working memory storage in the absence of considerable executive control requirements. Random sequences of digits were displayed on the screen, one digit every second and each digit displayed only once within a sequence. Following a sequence, as many question marks as digits were displayed on the screen and subjects were required to reproduce the digits either in the forward or backward order. The first sequence started with 3 digits, followed by sequences of increasing number of digits (up to 10) if the previous answer was correct. If not, a new sequence with the same number of digits was displayed. If the answer was incorrect again, the test stopped. The final score was the maximal number of digits that was answered correctly.

Finally, participants performed a computerized short version of the standard progressive matrices intelligence test using only the two most difficult matrix sets (D and E) out of all five sets ^62^. Each set comprised 12 matrices presented in progressively difficult order. The non-standardized intelligence score was the sum of correctly solved matrices.

### Experiment 2 specifics

Experiment 2 was designed as a follow-up to experiment 1 to specifically test the hypothesis that prefrontal cortex representations are confined to intentionally learned (here: instructed) S-R rules rather than incidentally learned S-R rules and further that these representations are not merely related to the identities of the involved stimuli or responses. Therefore, experiment 2 included the easy condition only (i.e. 4 instructed and implemented S-R rules per task) and two types of conditions were realized, including an instructed learning condition (as in experiment 1) and a control condition. Different from the learning condition, in the control condition the instruction cues (the vertical bars) were omitted during the instruction phase but were instead presented together with the nouns during the implementation phase (see Figure 2). Hence, in the control condition no S-R rules could be memorized during the instruction phase and task implementation could rely entirely on the explicit response cues rather than memorized instructions. Additionally, experiment 2 was designed to track the representational dynamics across a more extended practice period. Therefore, each noun was presented 8 times during the implementation phase (instead of 4 times in experiment 1). Measurements were taken in three consecutive runs (18 min each) comprising 4 blocks of each condition (instructed and control) in pseudo-randomized order, amounting to a total of 12 blocks per condition (total duration approximately 54 minutes). Before entering the scanner each participant completed a short practice session comprised of one task block for each condition with a separate stimulus set not used during the main experiment. Different from experiment 1, measures of working memory span and general intelligence were not taken.

### Behavioral data analysis

Behavioral performance was assessed regarding mean response times for correct responses (RTs) and regarding response accuracies (proportion of correct responses). Mean RTs and response accuracies were each analyzed with repeated measures ANOVAs. In experiment 1, response accuracies were especially relevant as a measure of representational integrity which was targeted by the manipulation of instruction difficulty. Since there was no feedback provided after response execution, representational integrity might be quantified inadequately if accuracy was measured in ‘objective’ terms with reference to the originally instructed response. The reason is that - in case the originally instructed response is not properly recalled – participants might generate subjectively defined rule representations based on the response that was actually executed for a specific stimulus irrespective of whether this was the originally instructed response. To account for this, response accuracies were defined relative to the response that was executed upon the preceding occurrence of a specific stimulus. Since there is by definition no response execution prior to stimulus repetition 1, accuracy was in this case naturally defined relative to the instructed response, thus providing an ‘objective’ accuracy measure. This definition of response accuracies was applied in both experiments.

### Imaging methods

#### Data acquisition

MRI data were acquired on a Siemens 3T whole body Trio System (Erlangen, Germany) with a 32 channel head coil. Ear plugs dampened scanner noise. After the experimental session structural images were acquired using a T1-weighted sequence (TR = 1900 ms, TE = 2.26 ms, TI = 900 ms, flip = 9°) with a resolution of 1 mm × 1 mm ×1 mm. Functional images were acquired using a gradient echo planar sequence (TR = 2000 ms, TE = 30 ms, flip angle = 80°). Each volume contained 32 slices that were measured in ascending order. The voxel size was 4 mm × 4mm × 4 mm (gap: 20%). In addition, field maps were acquired with the same spatial resolution as the functional images in order to correct for inhomogeneity in the static magnetic field (TR = 352 ms, short TE = 5.32 ms, long TE = 7.78 ms, flip angle = 40°). The experiment was controlled by E-Prime 2.0.

### Preprocessing

The acquired fMRI data were analyzed using SPM12 running on MATLAB R2016a. First, the functional images were slice-time corrected, spatially realigned and unwarped using the acquired field maps. Each participant’s structural image was co-registered to the mean functional image and segmented. Spatial normalization to MNI space was performed by applying the deformation fields generated by the segmentation process to the functional images (resolution: 3 mm × 3 mm × 3 mm). The images were not additionally smoothed prior to GLM estimation in order to suit the planned MVPA ^63^. Instead each subjects’ images were smoothed with 6 mm FWHM after the MVPA was completed.

### Voxelwise single-trial BOLD estimation

Voxel-wise BOLD activation was estimated based on the General Linear Model (GLM) approach implemented within the SPM12 framework using a first-order auto-regressive model and including a 1/128 Hz high-pass filter in experiment 1 and a 1/256 Hz high-pass filter in experiment 2 in order to accommodate different learning block lengths. During GLM estimation SPM’s implicit analysis threshold was switched off and instead all non-brain voxels were masked out using SPM’s intracerebral volume mask ‘mask_ICV.nii’. This procedure was chosen to enable group level statistics for regions affected by susceptibility-induced signal loss in a few subjects.

BOLD activations during the implementation phase were modeled by using single-trial GLMs. We used the least-squares-separate (LSS) model approach ^64,65^ which included one regressor modeling one specific trial and another regressor modelling all other trials, plus a constant. To obtain estimates for each single trial, we estimated as many different LSS models as there were trials. While LSS modeling is computationally much more time consuming, it has been argued to produce more robust estimates than other approaches ^64,65^. Regressors were created by convolving stick functions synchronized to stimulus onset with the SPM12 default canonical HRF. In experiment 1, this implies a total of 192 single-trial regressors per run (16 trials per task block times 12 task blocks), which amounts to 576 across all three runs. In experiment 2, this implies a total of 256 single-trial regressors per run (32 trials per task block times 8 task blocks), which amounts to 768 across all three runs.

In addition to the single trial regressors used for the implementation phase, we included regressors for the instruction phase and for the performance feedback at the end of each implementation phase. To appropriately capture BOLD activation during the instruction phase, spanning either 12 s (easy condition) or 14 s (difficult condition), we used Fourier basis set regressors including 20 different sine-wave regressors spanning 44 s which were time-locked to the onset of the start of the instruction phase. Using a Fourier basis set has the advantage to flexibly model any BOLD response shape associated with the extended instruction phase without making prior shape assumptions. An advantage over FIR modeling is that a Fourier basis set easily operates at micro-time resolution (SPM default TR/16) whereas FIR operates at TR resolution only ^66^. Performance feedback was modeled with a standard event-related HRF function time-locked to the onset of the feedback screen.

### Multivariate pattern analysis

The MVPA was based on single-trial beta estimates obtained for the implementation phase. Rule identity-specific activation patterns were determined by adopting a modified versions of the multi-voxel pattern correlation approach ^67,68^ geared towards the unbiased computation of time-resolved pattern correlations within runs. Specifically, identity-specific patterns were identified by computing the mean difference between (i) pattern correlations for re-occurrences of same stimuli and (ii) pattern correlations for occurrences of different stimuli. Such mean difference values were computed separately for each task block and within each task block separately for each successive learning stage defined by two consecutive occurrences per stimulus (Fig. 3).

This procedure allowed us to analyze two learning stages in experiment 1 (stage 1: stimulus repetitions 1 and 2; stage 2: stimulus repetitions 3 and 4). In experiment 2, two additional learning stages could be analyzed involving stimulus repetitions 5 and 6 (stage 3) and stimulus repetitions 7 and 8 (stage 4). Finally, for each subject, the resulting mean difference values were averaged across task blocks separately for each learning stage before being submitted to group-level statistical evaluation. Based on previous work, trial sequences were constructed in a way that ensured unbiased multivariate results under conditions of overlapping single-trial BOLD responses within task blocks ^15,64,69^. Specifically, we conducted unbiased identity-specific MVPA separately for each successive learning stage within each task block. To this end, for each task block the overall trial sequence was composed of 2 (experiment 1) or 4 (experiment 2) independently generated ‘atomic’ 8-trial sequences, each comprising 2 randomly distributed occurrences of each of the 4 nouns. On average across such atomic sequences, this approach guaranties unbiased MVPA due to the circumstance that non-zero bias regarding individual atomic sequences is distributed around zero mean ^15^. We furthermore took advantage of multiple novel task blocks per participant which allowed us to regress out bias-induced variance across blocks and thereby to obtain more robust results. Bias-induced variance regarding pattern similarity estimates was determined for subject-specific white-matter volumes (see below) with verified absence of significant multivariate effects on average across subjects (see Figure 6B).

The primary MVPAs were computed for regions-of-interest (ROIs) considering all voxels within a ROI simultaneously. Additional searchlight-based MVPAs were computed for the whole-brain volume with a spherical searchlight radius of 3 voxels ^63^ as implemented in the CosmoMVPA toolbox ^70^. The ROI-based approach was employed to be able to conveniently compare multivariate effects between different LPFC regions in a proper statistical way ^39^. The complementary searchlight approach allowed us to also identify additional effects outside the pre-specified ROIs. Additionally, the searchlight approach was used to localize MVPA effects within the anatomical ROIs with better spatial precision. Searchlight results were statistically evaluated at the peak-level with p<.05, FWE-corrected for the whole-brain volume or for the ROI volume, respectively.

Four anatomically constrained ROIs were included based on the previous literature which had most consistently highlighted the potential relevance of lateral PFC regions ^28,29,31,34,35,43^. Using the automatic anatomic labeling atlas ^71^, we included for each hemisphere the ventrolateral PFC (according to the combined aal regions ‘inferior frontal gyrus pars opercularis’ and ‘inferior frontal gyrus pars triangularis’) and the dorsolateral PFC (aal region ‘middle frontal gyrus’).

The MVPA was based on all trials including correct trials and error trials alike. This allowed us to test how differences in the overall proportion of error trials would modulate the strength of identity-specific pattern similarity effects. This procedure was preferred over running separate MVPAs selectively based on either correct trials or error trials. Especially, the relatively small proportion of error trials in the easy instruction condition renders reliable pattern similarity estimates unfeasible.

### Univariate analysis of mean activity

The MVPA was complemented by a conventional univariate analysis computed for the MVPA ROIs. Instead of single-trial beta estimates, the univariate analysis was based on beta estimates collapsed across all trials per condition. In experiment 1 the conditions were defined by easy blocks vs. difficult blocks and by stimulus repetitions (1 to 4). In experiment 2 the conditions were defined by instructed blocks vs. control blocks and by stimulus repetitions (1 to 8). Error trials were excluded.

### Functional connectivity analysis

Functional connectivity changes were computed specifically for experiment 2 using the beta-series correlation approach based on the same single-trial estimates that were already generated for the MVPA ^72-74^. Error trials were excluded. Following-up on previous study results ^20,24^, we examined functional connectivity changes comparing late learning trials (stimulus repetitions 7 and 8) with early learning trials (stimulus repetitions 1 and 2) with a special focus on connectivity between the lateral PFC and the basal ganglia.

## Acknowledgements

This work was funded by the German Research Foundation (Deutsche Forschungsgemeinschaft, DFG), CRC 940, subprojects Z2 and A2.

## Author Contributions

Conceptualization: H.R., U.W., T.S.; Methodology: H.R., H.M., T.S.; Formal Analysis: H.R.; Interpretation: H.R., T.S., K.Z., H.M., U.W.; Writing – Original Draft: H.R.; Writing – Review & Editing: H.R., T.S., K.Z., H.M., U.W.

